# Exclusive D-lactate-isomer production during a reactor-microbiome conversion of lactose-rich waste by controlling pH and temperature

**DOI:** 10.1101/2023.10.11.561877

**Authors:** Dorothea M. Schütterle, Richard Hegner, Monika Temovska, Andrés E. Ortiz-Ardila, Largus T. Angenent

## Abstract

Lactate is among the top-ten-biobased products. It occurs naturally as D– or L-isomer and as a racemic mixture (DL-lactate). Generally, lactate with a high optical purity is more valuable. In searching for suitable renewable feedstocks for lactate production, unutilized organic waste streams are increasingly coming into focus. Here, we investigated acid whey, which is a lactose-rich byproduct of yogurt production, that represents a considerable environmental footprint for the dairy industry. We investigated the steering of the lactate-isomer composition in a continuous and open culture system (HRT = 0.6 d) at different pH values (pH 5.0 *vs.* pH 6.5) and process temperatures (38°C to 50°C). The process startup was achieved by autoinoculation. At a pH of 5.0 and a temperature of 47°C-50°C, exclusive D-lactate production occurred because of the dominance of *Lactobacillus* spp. (> 95% of relative abundance). The highest volumetric D-lactate production rate of 722 ± 94.6 mmol C L^-1^ d^-1^ (0.90 ± 0.12 g L^-1^ h^-1^), yielding 0.93 ± 0.15 mmol C mmol C^-1^, was achieved at a pH of 5.0 and a temperature of 44°C (*n* = 18). At a pH of 6.5 and a temperature of 44°C, we found a mixture of DL-lactate (average D-to-L-lactate production rate ratio of 1.69 ± 0.90), which correlated with a high abundance of *Streptococcus* spp. and *Enterococcus* spp. However, exclusive L-lactate production could not be achieved. Our results show that for the continuous conversion of lactose-rich dairy waste streams, the pH was a critical process parameter to control the yield of lactate isomers by influencing the composition of the microbiota. In contrast, temperature adjustments allowed the improvement of bioprocess kinetics.

## 1. Introduction

The transition from a linear economy to a circular economy requires the replacement of fossil resources for the production of chemical building blocks and fuels. Organic carbon sources from agriculture can also be utilized as bio-based renewable resources (Tan and Lamers 2021). This poses problems when these renewable resources compete with food production. Therefore, exploiting organic waste streams has increasingly become the focus of attention. For the dairy industry, acid whey, which is a waste stream of strained yogurt production, presents a considerable environmental footprint due to large volumes and high chemical oxygen demand (COD) concentration of > 70 g L^-1^ (Sar et al. 2022, Xu et al. 2018). In contrast to the production of food nutrients, such as whey powder from sweet whey (Atra et al. 2005), the dairy industry is searching for economically and environmentally sustainable acid-whey-derived products (Rocha-Mendoza et al. 2021). It is challenging to dry acid whey to powder due to the acidic pH of approximately 4.4 (Miller et al. 2006), high levels of calcium, and a relatively high content of lactate (Chandrapala et al. 2015, Chandrapala et al. 2017).

Lactate is ranked as one of the top-ten-biobased products from biorefineries by the US Department of Energy (Bozell and Petersen 2010). The global lactate market is driven by the food, cosmetics, and pharmaceutical industry and demands around 1.22 million tons per year (Alves de Oliveira et al. 2018). One pertinent lactate demand is for the production of biodegradable poly-lactic acid (PLA) (Castro-Aguirre et al. 2016, Klotz et al. 2017). Future increasing demands for lactate are being anticipated. For example, lactate can be used for the fermentative production of medium-chain-carboxylates (MCCs) (Kucek et al. 2016, Zhu et al. 2015). MCCs have a higher economic value and are easier to separate from solution than lactate (Dahiya et al. 2018, Perimenis et al. 2018), and can be sold as sustainable antimicrobial feed additives or precursor molecules for biofuels and biochemicals (Agler et al. 2011, Solis de los Santos et al. 2009).

Lactate exists in three forms: D-lactate, L-lactate, and a racemic mixture (*i.e.*, DL-lactate) (Huang et al. 2021). A racemic mixture can be obtained through chemical synthesis, whereas the industrial route for optically pure D– or L-lactate production is through microbial fermentation with lactic acid bacteria (LAB) (Rawoof et al. 2020). Optically pure lactate isomers are considered more valuable than racemic mixtures (Huang et al. 2021). The fermentation process for L-lactate production has been industrially established since 1891 (Benninga 1990). *Streptococcus* spp., such as *Streptococcus thermophilus* St20, produce mostly L-lactate when cultivated in milk (Panesar et al. 2007). *Enterococcus* spp., such as *E. gallinarum* and *E. casseliflavus*, also produce L-lactate and are an essential part of the dairy industry (Giraffa et al. 2022). *Enterococcus faecium* S.156 produces L-lactate industrially at temperatures from 32°C to 40°C and pH values from 5.5 to 6.5 (Sun et al. 2015). On the other hand, LAB that are capable of producing D-lactate in adequate purity are relatively rare (Klotz et al. 2016). One D-lactate-producing species is *Lactobacillus delbrueckii* ssp. *bulgaricus*, which is a well-known yogurt bacterium (Ben-Yahia et al. 2012).

In the last two decades, D-lactate has gained considerable importance (Klotz et al. 2016). For example, an ideal optical isomeric (*i.e.*, enantiomeric) ratio of D-/L-lactate is crucial for the thermal and mechanical properties of PLA (Alexandri et al. 2019). Furthermore, D-lactate is the preferred enantiomer for MCC production by *Megasphaera elsdenii* due to the direct conversion of D-lactate to pyruvate, while L-lactate conversion requires lactate racemase (Hino and Kuroda 1993). Thus, lactate-based chain elongation would prefer D-lactate as the substrate compared to L-lactate (Kucek et al. 2016), especially because the competing pathway of lactate-based chain elongation, which is known as the acrylamide pathway, utilizes L-lactate.

Conventional microbial lactate-fermentation systems constitute about 40%-70% of the entire costs for sugars as substrates (Rawoof et al. 2020). It is, therefore, economically attractive to utilize organic waste streams. Acid whey is a carbohydrate-rich waste stream containing predominantly lactose, which is readily available for microbial fermentation without any physical or enzymatic pretreatment. Several studies have already shown lactate production from different types of whey streams with pure cultures. An engineered acid-tolerant *Saccharomyces cerevisiae* has been used to convert lactose from sterilized acid whey into lactate during a batch fermentation (Turner et al. 2017). For a different study, *Lactobacillus casei* was used to ferment reconstituted and sterilized whey powder to L-lactate in a batch process (Panesar et al. 2010). Finally, immobilized *L. casei* was also studied continuously to ferment sterilized whey permeate into L-lactate (Krischke et al. 1991).

However, the conversion of acid-whey waste into lactate by pure-culture fermentation requires sterile process conditions, which is costly. Conversely, open-culture biotechnology using reactor microbiomes offers distinct advantages compared to pure-culture fermentation. Due to their metabolic diversity, reactor microbiomes have proven to be stable biocatalysts for continuous processes, allowing complex organic waste streams to be processed efficiently without sterilization (Kim et al. 2016, Kucek et al. 2016). Because LAB are used in yogurt production, we theorized that the indigenous microbes in acid whey could be enriched within reactor microbiomes to convert the left-over lactose into lactate. Commonly used dairy LAB produce either D-, L-, or DL-lactate, and exhibit distinct growth and production optima based on pH and temperature (Adamberg 2003, Panesar et al. 2007). Microbiome studies with food waste have, indeed, shown different lactate production characteristics by changing the temperature (Daly et al. 2020). Here, we examined the effects of pH and temperature on the lactate-isomer production spectra from unsterile acid-whey waste in a continuously operated system with a reactor microbiome.

## 2. Material and methods

### 2.1 Acid whey and autoinoculation

We utilized undiluted acid whey without adding vitamins, minerals, yeast extract, and carbon sources. FrieslandCampina Germany GmbH provided the acid-whey batch, which we used throughout the study, from their production site in Cologne, Germany on September 21, 2021. The acid whey was transferred in a 300-L intermediate bulk container from Cologne to our laboratories at the University of Tübingen. Upon delivery, liquid samples for biological and chemical analysis were taken (**Table S1**), and the acid-whey batch was stored in 5-L aliquots at –20°C until further use. *Prior* to use, the acid whey was thawed in a water bath and transferred to a 2-L feed tank (DWK life-sciences, Mainz, Germany). The acid whey in the feed tank was continuously mixed during the experimental period, using a magnetic stir plate (Stuart® US151, Cole-Parmer, Vernon Hills, IL, USA). The acid-whey feed tank was kept refrigerated in a 10-L ice-filled plastic bucket that was insulated while being open to the atmosphere. Samples were taken regularly and analyzed to determine the quality (**Table S2**). The bioreactor operating experiment was started by autoinoculation, and thus no external inoculum was added.

### 2.2 Bioreactor system

The experimental setup consisted of four glass-jacketed bioreactors (**Fig. S1**), with working volumes of 250 ml (**Table S3**), which were operated in parallel as duplicates at a pH of 5.0 or 6.5. We controlled the temperature using a recirculating thermostatic water bath (Huber, Offenburg, Germany). Each bioreactor was closed with a 3-D printed lid made of PLA (Innofil3D, the Netherlands) and a rubber-mason-jar seal. We equipped each bioreactor with a pH electrode and a temperature sensor (Avantor, Radnor, PA), which were connected to a pH controller (Bluelab, Gisborne, New Zealand) and a 2 M NaOH solution. We used a multichannel pump (Cole-Parmer GmBH, Wertheim, Germany) to continuously feed acid whey and remove fermentation broth to achieve an HRT of 0.6 d. All bioreactors were sparged with dinitrogen gas throughout the operating period to maintain anaerobic conditions. They were continuously mixed at 250 rpm using a magnetic multi-stirrer plate (2mag AG magnetic emotion, Munich, Germany).

### 2.3 Experimental Design

The experiment was split into three separate operating periods, which we refer to as P-I, P-II, and P-III (**Table 1**). After a startup period of 4 days in batch mode at 44°C (left grey area in **Fig. 1**), the bioreactors were operated in continuous mode with a temporary change to batch mode between P-I, P-II, and P-III (four grey areas in **Fig. 1**). We gradually increased the temperature from 44°C to 50°C between Day 4 and 8 (**Fig. 1A**). Next, we lowered the temperature after every third day (*i.e.*, after 72 h) in steps of 3°C to obtain a temperature gradient from 50°C to 38°C during P-I (**Fig. 1A**). The parameters for P-II and P-III were chosen based on the results of P-I.

**Figure 1.**
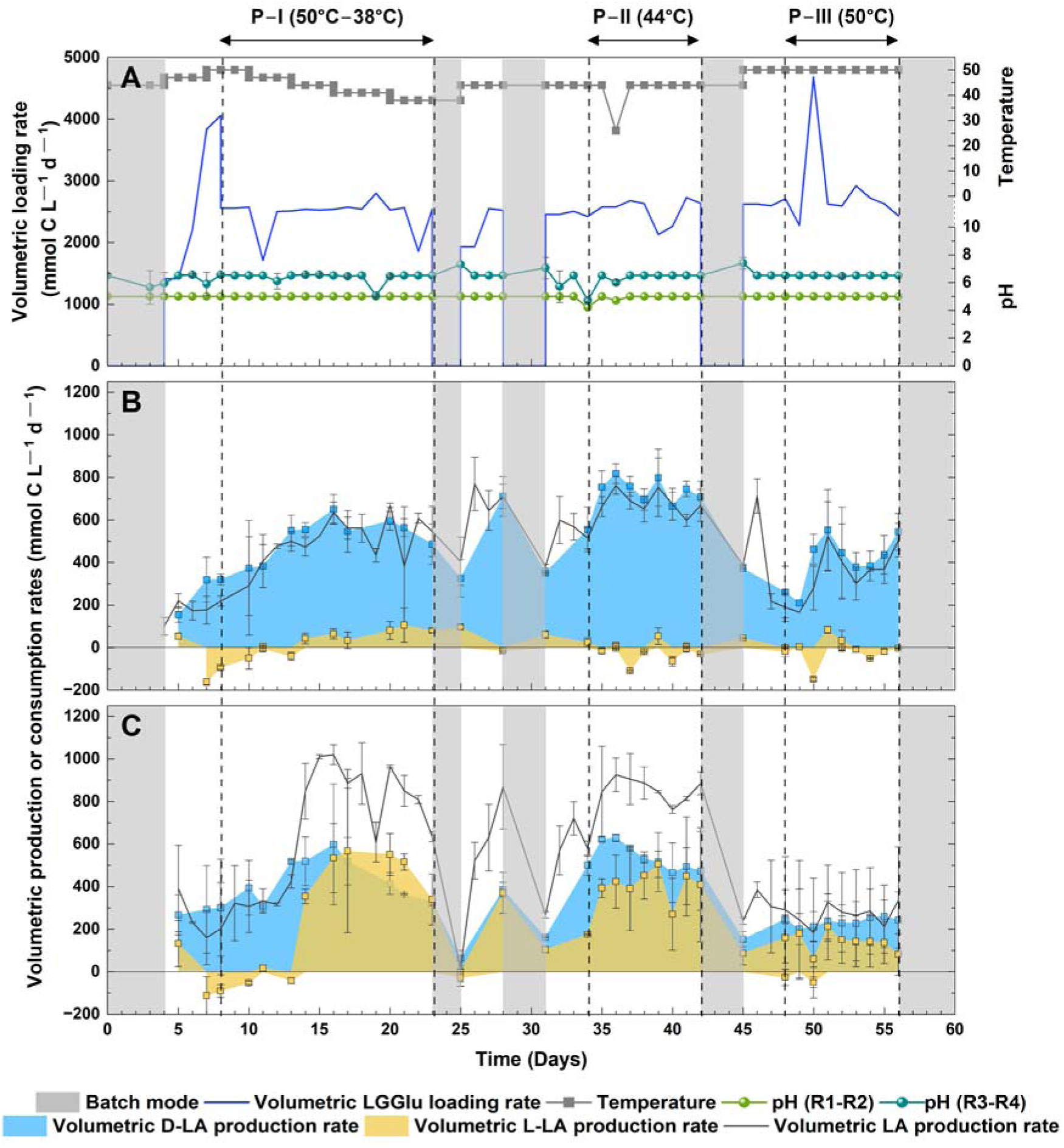
Performance of the reactors throughout the operating period (P1-PIII): **A)** line plots of the volumetric LGGlu loading rate (lactose, galactose, and glucose, excluding the lactate present in the acid whey), temperature, and pH; **B)** line plots of the total volumetric lactate production rate and area plots of the volumetric D-lactate and L-lactate production and/or consumption rates for reactor duplicates at pH 5; and **C)** line plots of the total volumetric lactate production rate and area plots of the volumetric D-lactate and L-lactate production and/or consumption rates for reactor duplicates at pH 6.5. Error bars represent the standard deviations among duplicate reactors.

**Table 1.** Process parameters during operating periods P-I, P-II, and P-III.

| Parameter | Period I (P-I) | Period II (P-II) | Period III (P-III) |
| --- | --- | --- | --- |
| Duration | Day 8 to 23 | Day 34 to 42 | Day 48 to 56 |
| Temperature | 50°C – 38°C | 44°C | 50°C |
| Mode of operation | Continuous | Continuous | Continuous |

### 2.4 Analyses

We collected 1 mL of liquid samples during P-I, P-II, and P-III on a daily basis and prepared them for storage (Esquivel-Elizondo et al. 2021). The concentrations of glucose (Glu), lactose (L), galactose (G), lactate (LA, the total of the dissociated and undissociated form), and ethanol were measured by HPLC (LC20, Shimadzu, Japan), which was equipped with an Aminex HPX-87H column and a refractive index detector (RID) (Klask et al. 2020). The concentrations of D– and L-lactate isomers were measured in technical duplicates with the D-/L-lactic acid kit (Megazyme, Bray, Ireland). We measured the carboxylates acetate, propionate, *n*-butyrate, *n*-valerate, *n*-caproate, and *n*-caprylate by an Agilent 7890B GC (Agilent Technologies, Inc., Santa Clara, CA, United States), which was equipped with a capillary column (DB-Fatwax UI 30 m × 0.25 m; Agilent Technologies, Inc., Santa Clara, CA, United States) and an FID detector (Esquivel-Elizondo et al. 2021). We made the following modifications for the instrument method and sample preparation: The injection and detector temperatures were set to 200°C and 250°C, respectively. Ethyl-L-Lactate was used as an internal standard, and 2% formic acid was used for sample acidification (to pH 2).

For volatile suspended solids (VSS) measurements, fermentation broth samples were taken on Days 10, 13, 16, 20, 28, 35, 42, 49, and 56 and stored in 15-mL falcon tubes at –20°C until analysis. VSS were measured according to the Standard Methods For the Examination of Water and Wastewater (Sections 2540 D and E) (Baird et al. 2012) with the following modifications: Volumes between 7 mL to 10 ml were used to reach a weight of min 2.5 mg to max. 200 mg of dried residue on the filter. The glass microfiber filters (Whatman, Maidstone, Great Britain) were dried overnight at 80°C and under vacuum. After and before weighing, residues on glass microfiber filters were ignited in a muffle furnace (L 3/12/B410, Nabertherm, Lilienthal, Germany) at 550°C for 30 min.

### 2.5 Microbial community analysis

#### 2.5.1 Biomass sampling, DNA extraction, 16S-rRNA-gene sequencing, and data analysis

We monitored the reactor microbiome throughout the operating periods by 16S rRNA gene sequencing. During P-I, samples were taken on Day 10, 13, 16, and 20. To assess the final reactor microbiome compositions after P-II and P-III, samples were taken on Day 42 and 56, respectively. We took 1.5 mL of liquid broth from each reactor. Next, these samples were centrifuged for 6 min at 16,740 x g at room temperature (5427 R, Eppendorf, Hamburg, Germany), and the supernatant was discarded. The cell pellets were stored at –80°C in sterile reaction vials with screw caps (Carl Roth GmbH, Karlsruhe, Germany) until further processing.

DNA isolation was performed with the FastDNA SPIN Kit for Soil (MP Biomedicals, Solon, OH). The DNA integrity was assessed by 1% agarose Midori Green Advanced stained gel electrophoresis, according to Cheng et al. (2013). 16S rRNA gene amplification of the V4 region was accomplished by polymerase chain reaction (PCR) using the universal 515F and 806R primers (IDT, Coralville, IA) with Illumina overhang sequences (**Table S4**). PCR products were cleaned using AMPure XP magnetic beads (Beckman Coulter, Brea, CA) until a clear, unique band was detected by 1% agarose gel electrophoresis. MiSeq Illumina library preparation was performed using barcoded DNA with the Nextera XT Index kit V2 (Illumina, San Diego, CA), following the manufacturer’s instructions for dual indexing. Genomic library products were quantified with a dsDNA Qubit spectrophotometer (Qubit Flex Fluorometer, Thermo Fisher Scientific, Singapore), standardized at a 4-nM concentration, and pooled for the MiSeq Illumina platform. Sequencing was carried out using the MiSeq Reagent Kit v2 (300 cycles) (**Fig. S2**). The raw fastq files were deposited at the National Center for Biotechnology Information (NCBI) database (Accession number: PRJNA885033).

#### 2.5.2 Sequencing data processing

From the obtained sequences, 16S rRNA gene amplicons were processed *via* Quantitative Insights Into Microbial Ecology 2 (QIIME2 – v 2021.8.0) (Bolyen et al. 2019). Sequences were demultiplexed using the QIIME default pipeline. Quality filtering, sequence joining, chimera removal, and general denoising were performed using the Divisive Amplicon Denoising Algorithm (DADA2) (Callahan et al. 2016). Obtained representative sequences were filtered, singletons and low-frequency sequences were removed, and then taxonomically classified using machine learning with the Scikit-learn naive-Bayes custom-trained classifier (Pedregosa et al. 2012). Greengenes 16S rRNA gene database version 13_8 was used to train the naive-Bayes classifier with the primers (*i.e.,* 515F – 806R), setting a 75% acceptance as a cut-off match identity with the filtered representative sequences.

We estimated α diversity (*i.e.*, microbial richness and abundance for each sample) and β diversity (*i.e.*, comparisons between samples) (Werner et al. 2012). The α diversity was evaluated using the QIIME2 diversity α-rarefaction pipeline with Shannon and dominance indexes (Johnson and Burnet 2016). To test for significant differences in α diversity between pH conditions and applied temperatures, the Kruskal-Wallis tests (pairwise) were used. The β diversity was evaluated using the Bray-Curtis dissimilarity index and UniFrac diversity tests (Lozupone and Knight 2005). We used principal coordinate analysis (PCoA) to visualize the β diversity turnover using a Bray-Curtis distance matrix. We performed a distance-based constrained redundancy analysis (dbRDA) (Legendre and Anderson 1999) with operative variables as constraints (*i.e.,* yields, consumption rates), correlating the change in the microbial community structure with the operative conditions as eigenvectors (Agler et al. 2012). Finally, a Direct Gradient Analysis (DGA) (ter Braak 1986) was used to ordinate the most significant microbial communities as eigenvectors (ANOVA, *p* < 0.05, relative abundance ≥ 1%) with the constraints used in the dbRDA analysis. dbRDA, DGA, and plots were performed with the Vegan community ecology package for R (Oksanen et al. 2012). Relative abundance plots were generated in OriginPro, Version 2023 (OriginLab Corporation, Northampton, MA, USA.), and PCoA with dbRDA plots were made in RStudio *via* the ggplot2 package (Wickham 2011).

### 2.6 Calculations and statistical analysis

We reported the concentration of carbon-containing compounds as mmol C L^-1^ and calculated the volumetric loading rates based on the HRT. Consequently, product yields are given as mmol C_product_ mmol C^-1^_consumed substrate_ (supporting information [SI]; **Eq. S1-S4**). The volumetric lactate (isomer) production rate refers to the net lactate (isomer) production rate within the bioreactor system and has been corrected for the influent lactate (isomers) feeding rate, which is inherent to feeding acid whey. After each temperature change, the system was given an adaptation time of at least 3 HRTs (*i.e.*, 1.8 days) for all operating periods to reach steady-state conditions. Therefore, all values are generally given for a point in time/period after an adaptation period of at least two full days after the start of the operating period. Descriptive statistics (mean, standard deviation [SD], median, and skewness) of the kinetic bioprocess performance parameters (**Table S5, S6**) were analyzed in OriginPro, Version 2023. We refered to only the mean ± SD in the main text when the skewness was between –1 and 1. When the skewness was out of this range, we reported both the mean and the median by the mean (median) ± SD in the text.

For the comparative statistical analyses, the parametric statistical assumptions were tested using the Levenne (homoscedasticity), Kolmogorov-Smirnov (normality), and residual regression model (graphical normality) tests. Then, the data were analyzed under a General Linear Model (GLM) using the factorial analysis of variance (2-way ANOVA) test to elucidate the interactions between the operating variables and the D– and L-lactate isomers. After the ANOVA tests, the HDS Tukey and Scheffé tests were used as *post-hoc* analyses to define the pairwise differences between treatments. In the case of not acceptance of the parametric statistics assumptions, non-parametric tests of Kruskal-Wallis to test the significant statistical differences and Dunn pairwise comparisons were used as *post-hoc* analyses. Finally, Pearson and Spearman’s correlations were used as a confirmatory analysis of the dependent relationship between the variables and the changes in microbial abundance. For all the tests, the α error was assumed to be 0.05 under a significance of 95%, using the IBM SPSS statistics software (IBM) and the Statistix 10 (AnalyticalSoftware).

## 3. Results and Discussion

### 3.1 A mildly acidic pH of 5.0 and temperature steps of 47°C and 50°C supported exclusive D-lactate production within a reactor microbiome

The startup of the bioreactor experiments *prior* to continuous operation during P-I was conducted by autoinoculation in batch mode for four days at 44°C (**Fig. 1A**). For the operation of the four bioreactors (duplicates), we chose two different pH values *(i.e.*, 5.0 and 6.5) and a temperature gradient (*i.e.*, 38°C to 50°C) in accordance with published data on the growth conditions of LAB (Adamberg 2003). A temperature gradient was utilized during P-I, in addition to the two different pH values (**Fig. 1A**), to also screen for an optimum temperature for the production of lactate isomers. For the two bioreactors with a controlled mildly acidic pH of 5.0, we observed an apparent production of primarily D-lactate throughout the entire temperature gradient during P-I (**Fig. 1B**). The yield of D-lactate was not significantly different throughout the entire temperature gradient between 38°C and 50°C at a pH of 5.0 (*p* = 0.078, HSD Tukey). Regardless, we observed exclusive D-lactate production only for the 47°C and 50°C temperature steps during P-I, except that very small amounts of L-lactate were produced for some samples at 47°C, resulting in a median of zero (**Table S5**). To our knowledge, this is the first report on exclusive D-lactate production with a reactor microbiome that was operated as an open and continuous culture.

The dominant lactate isomer in the acid whey was the L-lactate isomer *vs*. the D-lactate isomer, with an average concentration of 88.4 ± 17.7 mmol L^-1^ and 1.75 ± 1.79 mmol L^-1^, respectively (**Table S2**). Exclusive D-lactate production in the bioreactors meant that L-lactate remained in the effluent, albeit small amounts of L-lactate were consumed (**Fig. 1B, 1C** and **Table S5, S6**). We did not observe pertinent L-lactate consumption by lactate conversion or isomerization (Desguin et al. 2014). However, at temperatures between 38°C and 44°C during P-I, we detected some L-lactate production at a pH of 5.0 (**Fig. 1B** and **Table S5**). Thus, temperatures between 38°C and 44°C created a minor niche for L-lactate-producing bacteria next to D-lactate-producing bacteria. This minor niche was unavailable for the 47°C and 50°C temperature steps, and therefore the exclusivity for D-lactate production.

### 3.2 The volumetric D-lactate production rates were higher at 44°C than at 50°C for both pH values

During the temperature gradient study at a pH of 5.0, we observed that besides some L-lactate production, the total volumetric lactate production rate had increased at the lower temperatures to a maximum of 556 ± 95.2 mmol C L^-1^ d^-1^ (n = 8) at a temperature of 41°C (**Fig. 1B** and **Table S5)**. The step-wise temperature drop increased the volumetric D-lactate production rate to a maximum of 603 (589) ± 70.4 mmol C L^-1^ d^-1^ (n = 4) at a temperature of 44°C, while this was 347 (320) ± 133 mmol C L^-1^ d^-1^ (n = 4) at a temperature of 50°C and a pH of 5.0 (**Table S5**). We decided to further study the kinetic advantage at the lower temperatures by operating the four bioreactors for a more extended operating period at a temperature of 44°C (P-II) and 50°C (P-III). Indeed, our temperature gradient results during P-I were verified with a maximum volumetric D-lactate production rate of 722 ± 94.6 mmol C L^-1^ d^-1^ (n = 18) during P-II (pH 5.0 and 44°C) and a significantly lower volumetric D-lactate production rate of 420 ± 136.5 mmol C L^-1^ d^-1^ (n = 17) during P-III (pH 5.0 and 50°C) (**Table S6**) (*p* < 0.05, HSD-Tukey).

At the pH of 5.0, the D-lactate yields remained high with values of 0.93 ± 0.15 (n = 18) mmol C mmol C^-1^ at 44°C (P-II) and 0.97 ± 0.38 (n = 15) mmol C mmol C^-1^ at 50°C (P-III) (**Table S6**). For both temperatures, primarily D-lactate was produced (**Fig. 1B**), with L-lactate yields of 0.01 (0.00) ± 0.03 mmol C mmol C^-1^ at 44°C (P-II) (n = 18) and at 50°C (P-III) (n = 13) (**Table S6**). The high D-lactate yields throughout the entire operating period at a pH of 5.0 were in stark contrast with the two bioreactors that were operated at a pH of 6.5 (*p* < 0.05, HSD-Tukey). During P-I, the results for the initial two temperature steps of 47°C and 50°C were still almost identical for all bioreactors (**Fig. 1B,C** and **Table S5**). At temperatures between 38°C and 44°C during P-I, however, the volumetric L-lactate production rates became similar to the volumetric D-lactate production rates at a pH of 6.5 (**Fig. 1C** and **Table S5)**. This matched the L– and D-lactate yields (**Table S5**), which remained this way throughout the extended operating period at a temperature of 44°C (P-II) and 50°C (P-III) (**Table S6**). For example, the L-lactate yield was 0.22 ± 0.06 mmol C mmol C^-1^ (n = 18), while the D-lactate yield was 0.32 ± 0.10 mmol C mmol C^-1^ (n = 18) at 44°C (P-II) (**Table S6**).

Even though the bioreactors that were operated at a pH of 6.5 were producing a racemic mixture (DL-lactate) throughout the remainder of the operating period, the volumetric D-lactate production rate was higher at a temperature of 44°C than at 50°C and not even that much lower than in the bioreactors that were operated at a pH of 5.0. An average volumetric D-lactate production rate of 535 ± 106 (n = 18) mmol C L^-1^ d^-1^ and 235 ± 76.8 mmol C L^-1^ d^-1^ (n = 18) was achieved during P-II (pH 6.5 and 44°C) and during P-III (pH 6.5 and 50°C), respectively (**Table S6**). This resulted in a considerable and statistically higher total volumetric lactate production rate for the bioreactors that were operated at a pH of 6.5 than 5.0 at a temperature of 44°C during P-II (**Fig. 1B,C**) (*p* < 0.05, HSD-Tukey).

Thus, the temperature influenced the volumetric lactate production rate. For both pH levels that we tested, the volumetric D– and L-lactate production rates were higher at 44°C compared to 50°C. The highest volumetric isomeric lactate production rates that were reached here were 722 ± 94.6 (n = 18) mmol C L^-1^ d^-1^ or 0.9 g L ^-1^h^-1^ (pH 5.0 and 44°C) and 385 ± 144 (n = 18) mmol C L^-1^ d^-1^ or 0.5 g L ^-1^h^-1^ (pH 6.5 and 44°C), for D– and L-lactate production, respectively (**Table S6**). Thus far, studies on lactate-isomer production with microbiomes were only conducted in batch-mode cultivation and were mainly focused on L– lactate production. Tashiro et al. (2016) produced up to 1.1 g L^-1^ h^-1^ of 100% optically pure L-lactate from kitchen refuse with a microbiome at 50°C. In addition, Sakai and Ezaki (2006) produced 1.4 g L^-1^ h^-1^ of lactate with an optical purity of L-lactate of 97% with a microbiome at 55°C.

### 3.3 The relative abundance of *Lactobacillus* spp. was positively correlated to the D– lactate yield

The D-lactate-producing population *Lactobacillus* spp. had a relative abundance of 6.19% in the freshly thawed acid-whey substrate (**Fig. 2A**). The L-lactate-producing population *Streptococcus* spp. was the most abundant population in these acid-whey samples with a relative abundance of 58.0% (**Fig. 2A**), explaining why acid whey contained mostly L-lactate (88 mmol L^-1^ L-lactate *vs*. 1.7 mmol L^-1^ D-lactate in **Table S2**). The residence time of acid-whey substrate in the cooled feeding tank for the experimental setup was long enough to change the microbial composition with an increase in the relative abundance of *Lactobacillus* spp. from 6.19% to 10.3%. *Acetobacter* spp. was the dominant genus (77.0%), while the relative abundance of *Streptococcus* spp. reduced from 58.0% to 10.2% (**Fig. 2A**). During the operating period of the bioreactors: **(1)** the four-day autoinoculation event at a temperature of 44°C; **(2)** the four-day initial period at a continuous feed to increase the temperature to 50°C; and **(3)** the first two days of P-I at continuous feed (a total of ten days) (**Fig. 1A**), *Lactobacillus* spp. completely overtook the microbiome composition in the bioreactors that were operated at a pH of 5.0. This dominance remained through P-I, with relative abundances ranging between 96.7% and 98.8% (**Fig. 2B**).

**Figure 2.**
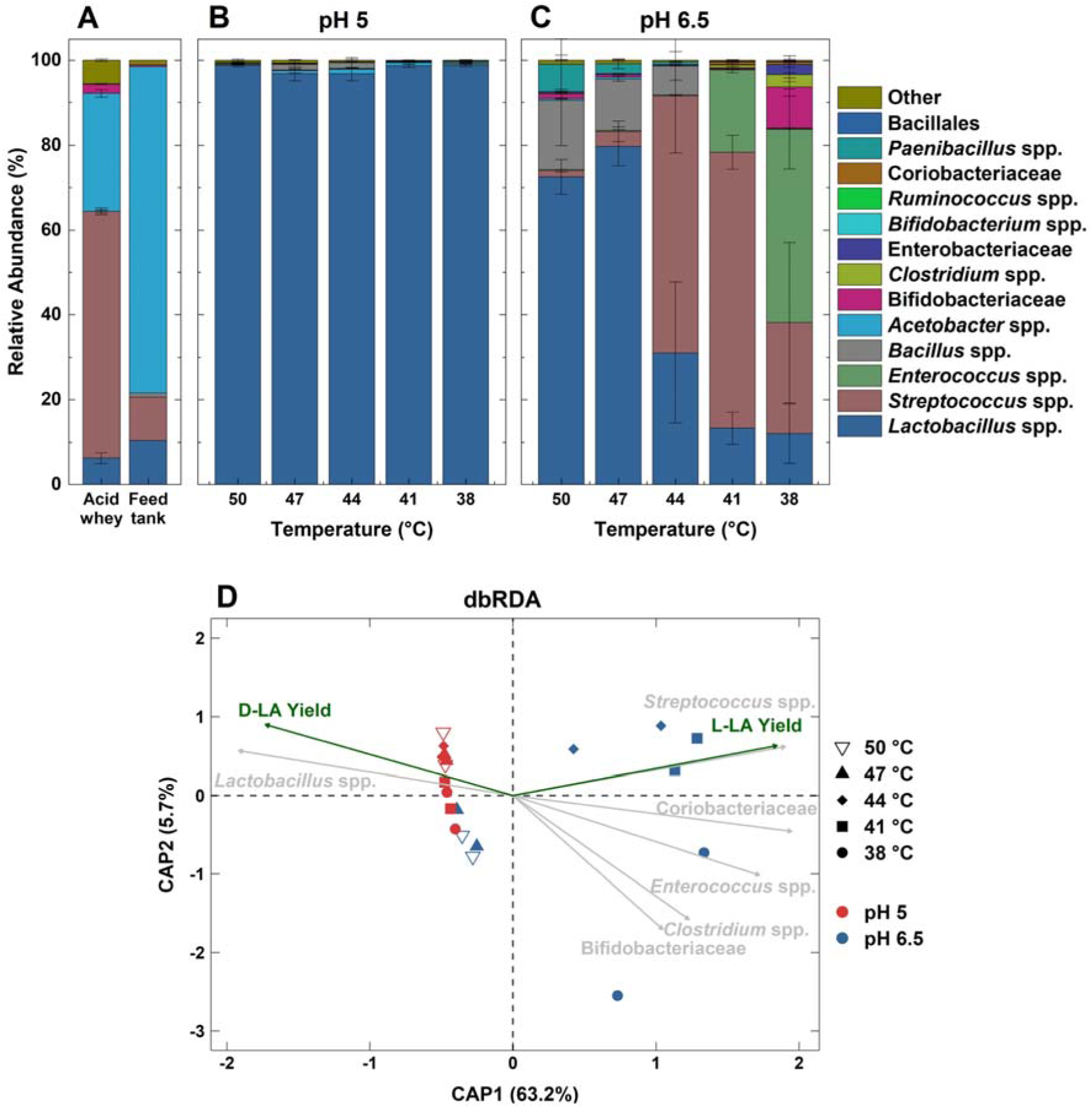
Microbial characterization during P-I: **A)** relative abundance of the microbial community for the freshly thawed acid whey (duplicate samples) and for the feed tank after one day (single sample); **B)** relative abundance of the microbial community at different temperatures throughout the operating period for duplicate reactors at pH 5.0; **C)** relative abundance of the microbial community for different temperatures throughout the operating period for duplicate reactors at pH 6.5; and **D)** distance-based redundancy analysis (dbRDA) of the microbial community and reactor parameters based on Bray-Curtis distance matrix. The percent variance explained by each axis is given in parentheses. Green arrows indicate functional performance parameters related to community composition. Grey arrows indicate populations that significantly influenced microbial community composition. Colors represent pH value and shapes represent applied temperature. Listed OTU’s comprised at least 1% relative abundance at the genus level. Error bars represent the standard deviation among duplicate reactors.

The dominance of *Lactobacillus* spp. in the two bioreactors that were operated at a pH of 5.0 was the reason for the exclusive D-lactate production finding during P-I (**Fig. 2B**). A further proof of a correlation between the relative abundance of *Lactobacillus* spp. and D-lactate yield was observed with the two bioreactors that were operated at a pH of 6.5 during P-I. Between 47°C and 50°C at a pH of 6.5, we measured predominant D-lactate production, which included L-lactate consumption, with almost similar results at the same temperatures at a pH of 5.0 (**Fig. 1B, 1C**). For these temperatures at a pH of 6.5, *Lactobacillus* spp. was again dominant, albeit less pronounced than at pH 5.0, with relative abundances between 72.5% and 79.8% (**Fig. 2C**).

When we then reduced the temperature from 47°C to 44°C during the temperature-gradient experiment at a pH of 6.5 (P-I), we observed a sudden emergence of L-lactate production with a significant increase in the L-lactate yield based on the HSD-Tukey’s test (*p* < 0.05) (**Table S5**). The volumetric D-lactate production rate increased as well, which we discussed above (**Fig. 1C**). *Lactobacillus* spp. was no longer dominant (31.1%), while the L-lactate-producing *Streptococcus* spp. became the dominant population with a relative abundance of 65.1% at 41°C (**Fig. 2C**). When we further decreased the temperature to 38°C, *Enterococcus* spp. became the most abundant population (45.5%). The loss of dominance in *Lactobacillus* spp. explained the shift in production from D-lactate towards a racemic mixture of DL-lactate (**Fig. 1C**, **Table S5**). Indeed, we observed a statistically relevant correlation between the relative abundance of *Lactobacillus* spp. and the D-lactate yield, which is visualized by the same direction for the arrows of *Lactobacillus* spp. and D-lactate yield (**Fig. 2D)**.

During our subsequent experimental periods (P-II and P-III), we had the objective of studying: **(1)** the stability of the *Lactobacillus* spp. dominance and its resulting high D-lactate yield throughout a longer operating period at a pH of 5.0; and **(2)** a possible return to a high D-lactate yield through *Lactobacillus* spp. dominance at a temperature of 50°C and a pH of 6.5 during P-III, which we had observed at the very beginning of P-I at the temperature steps of 47°C and 50°C. For the first objective (studying a possible *Lactobacillu*s spp. stability at a pH of 5.0), D-lactate was exclusively produced during P-II (44°C), resulting in a D-lactate yield of 0.93 ± 0.15 mmol C mmol C^-1^ (n = 18) (**Table S6**), with *Lactobacillus* spp. being the dominant population (98.2%) (**Fig. 3A**). During P-III (50°C), the average D-lactate yield slightly increased to 0.97 ± 0.38 mmol C mmol C^-1^ (n = 15) (**Table S6**), but this was not statistically relevant (*p* > 0.05, HSD-Tuckey). *Lactobacillus* spp. remained dominant (97.3%) (**Fig. 3A**), resulting in the absence of L-lactate production during P-II and P-III (no statistical difference: *p* > 0.05, HSD-Tukey) (**Fig. 1B**). Thus, throughout P-II and P-III the dominance of *Lactobacillus* spp. was stable at a pH of 5.0 with only four other OTUs reaching a relative abundance ≥ 1% (**Fig. 3A**).

**Figure 3.**
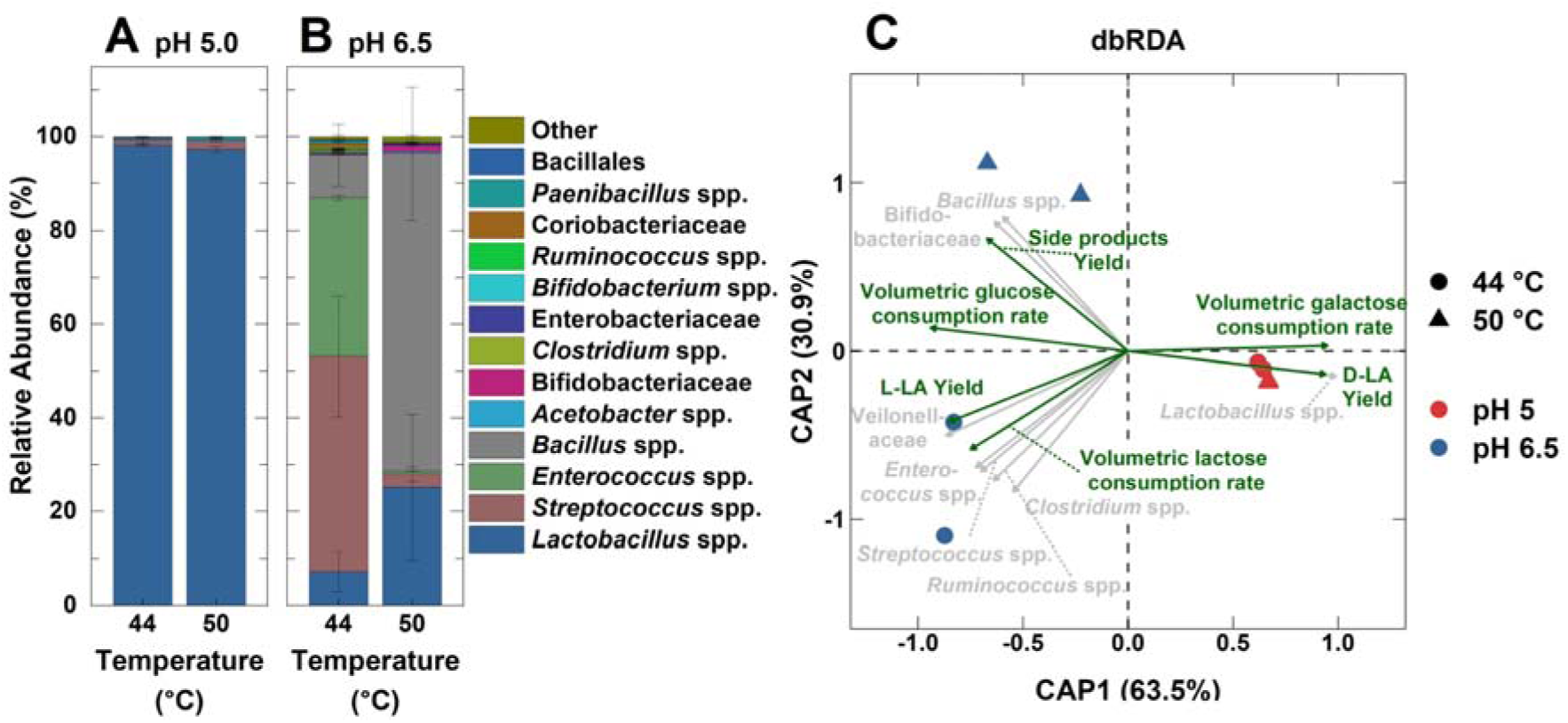
Microbial characterization during P-II and P-III: **A)** relative abundance of the microbial community at different temperatures throughout the operating period for duplicate reactors at pH 5.0; **B)** relative abundance of the microbial community for different temperatures throughout the operating period for duplicate reactors at pH 6.5; and **C**) distance-based redundancy analysis (dbRDA) of the microbial community and reactor parameters based on Bray-Curtis distance matrix. The percent variance explained by each axis is given in parentheses. Green arrows indicate functional performance parameters related to community composition. Grey arrows indicate populations that significantly influenced microbial community composition. Colors represent pH value and shapes represent applied temperature. Listed OTU’s comprised at least 1% relative abundance at the genus level, except for Veilonelaceae. Error bars represent the standard deviation among duplicate reactors.

Even though *Lactobacillus* spp. was dominant at a higher temperature of 50°C at a pH of 5.0, showing stability to temperature changes, the volumetric D-lactate production rates decreased. This likely indicates that the D-lactate-producing *Lactobacillus* spp. exhibits a sensitivity toward temperature. Adamberg et al. (Adamberg 2003) observed decreases in the growth yields and specific growth rates at temperatures > 47°C in *Lactobacillus bulgaricus* Lb12. Because we also measured a decrease in the VSS concentration in the bioreactors (**Fig. S5**), a lower growth rate of *Lactobacillus* spp. at a temperature of 50°C compared to 44°C explains the drop in the volumetric D-lactate production rates. In addition, dominant rod-shaped bacilli-like cell morphologies were observed during P-II at a pH of 5.0 (**Fig. S6**). The higher temperature of 50°C at a pH of 5.0 (**Fig. S6 C,D**), reduced the number of cells and elongated the rods compared to 44°C (**Fig. S6 A,B**), possibly identifying temperature stress.

The positive correlation between *Lactobacillus* spp. and D-lactate yield and its high level of stability at different temperatures at the mildly acidic pH value of 5.0, identified a specific ecological niche. For the L-lactate-producing *Streptococcus* spp., such as *S. thermophilus* St20, growth was only observed at pH values ≥ 5.1 (Panesar et al. 2007). In addition, Adamberg et al. (Adamberg 2003) showed in mono-culture and co-culture experiments that the growth rate of the D-lactate-producing *L. bulgaricus* Lb12 exceeded that of *S. thermophilus* St20 at pH ≤ 5.3. Thus, a mildly acidic environment (pH of 5.0) created a growth and competitive advantage for *Lactobacillus* spp. compared to *Streptococcus* spp. (De Angelis and Gobbetti 2004). In fact, *Lactobacillus* spp. seems to have evolved to thrive in mildly acidic conditions by not trying to maintain a constant internal pH during a pH decrease from 7.0 to 5.0 (Siegumfeldt et al. 2000), which is referred to as non-homeostasis. This relieves *Lactobacillus* spp. and other non-homeostatic LAB from too much stress from proton translocation, giving them a growth advantage. Indeed, the proton-translocating ATPase activity was the highest at mildly acidic conditions for some lactobacilli (De Angelis and Gobbetti 2004).

For the second objective (studying a possible return to a relatively high *Lactobacillu*s spp. abundance at a temperature of 50°C and a pH of 6.5), D– and L-lactate yields remained similar to each other at a temperature of 44°C (P-II), which was what we had anticipated (**Fig. 1C** and **Table S6**). L-lactate-producing *Streptococcus* spp. and *Enterococcus* spp. were the dominant population at this time point (45.9% and 34.0%, respectively), while the D-lactate-producing *Lactobacillus* spp. comprised 7.22% of the relative abundance (**Fig. 3B)**. Increasing the temperature to 50°C during P-III showed a small trend in shifting the lactate-isomer production toward D-lactate with a non-significant decrease in the L-lactate yield from 0.22 ± 0.06 mmol C mmol C^-1^ (n = 18) to 0.13 ± 0.10 mmol C mmol C^-1^ (n = 18). However, the D-lactate yield did not significantly change (*p* > 0.05, HSD-Tukey) (**Table S6**). The relative abundance of *Lactobacillus* spp. had increased from 7.22% to 25.2% after the switch from 44°C to 50°C, while that of *Streptococcus* spp. decreased. Still, we did not observe a switch to predominantly D-lactate production, which we had observed at very similar conditions (50°C at a pH of 6.5) at the beginning of the operating period (P-I) (**Fig. 1B**). *Lactobacillus* spp. did not reach the same relative abundances during P-III than during P-I (> 70%). Such hysteresis, for which more than one state can be observed under identical environmental conditions, is common in microbiome science (Allison and Martiny 2008).

### 3.4 How to operate a lactate-producing bioreactor to maximize value

The db-RDA analysis of the samples collected at P-II and P-III (*n* = 8) showed a positive correlation for the relative abundance of *Lactobacillus* spp. with D-lactate yield, while a positive correlation for the relative abundance of *Streptococcus* spp. and *Enterococcus* spp. with L-lactate yield was observed (**Fig. 3C**), confirming the results obtained during P-I (**Fig. 2D**). This statistical analysis also showed that the consumption of galactose, which is a breakdown product of lactose, correlated positively with *Lactobacillus* spp. abundance (**Fig. 3C**). Some strains of *L. bulgaricus* can utilize galactose *via* the Leloir pathway, where it is metabolized to lactate (Zourari et al. 1992), which would explain why a mildly acidic pH and high relative abundance of *Lactobacillus* spp. (**Fig. 2C**) coincided with galactose consumption. Galactose consumption is important to maximize product yield and to prevent unnecessary organic loads to the wastewater treatment plant at the biorefinery. For a pH of 6.5, the production, rather than the consumption, of galactose coincided with the high relative abundance of *Streptococcus* spp. and *Enterococcus* spp. (**Fig. 3B**). While only some strains of *S. thermophilus* are capable of galactose consumption, most strains release the galactose moiety back into the extracellular medium (Zourari et al. 1992).

Finally, the db-RDA analysis for the data of P-II and P-III identified that the relative abundance of *Bacillus* spp. and Bifidobacteriaceae was positively correlated to unwanted side products such as ethanol, acetate, propionate, and *n*-butyrate. The lowest side-product yield of 0.02 ± 0.01 mmol C mmol C^-1^ (n = 18) was measured at a pH of 5.0 and a temperature of 44°C during P-II (**Table S6**). This was associated with the dominance of *Lactobacillus* spp. (**Fig. 2B**) at mildly acidic pH and indicating that this population is homofermentative because little side product formation was detected. The temperature increase to 50°C during P-III at a pH of 5.0 resulted in a significant increase in the side-product yield to 0.11 (0.07) ± 0.09 mmol C mmol C^-1^ (n = 15) (HSD Tukey, *p* < 0.05) (**Table S6**), but the effluent concentrations remained relatively low (**Fig. S3C**). In contrast, these side-product concentrations became clearly elevated when the temperature was increased from 44°C to 50°C at a pH of 6.5 during P-III (**Fig. S4C**), resulting in the highest side-product yield of 0.15 (0.14) ± 0.05 mmol C mmol C^-1^ (n = 16) (**Table S6**). This coincided with a high relative abundance of *Bacillus* spp. (67.8%). Some *Bacillus* species, such as *Bacillus thermoamylovorans*, are heterofermentative and have been reported to produce ethanol an acetate, ethanol (Combet-Blanc et al. 1995), which we detected as well (**Fig. S4C**).

To maximize the value from acid-whey fermentation into lactate as a soluble intermediate product with a first-stage bioreactor for further processing with a second-stage bioreactor *(e.g.*, chain elongation), we would like to produce D-lactate at maximum yields. This means that: **(i)** galactose should not be wasted, and thus be consumed; and **(ii)** side-products should not be formed. Our work showed that the control of the pH was a decisive environmental parameter in tuning reactor microbiomes to perform different functions and that a pH of 5.0 *vs*. 6.5 showed a significant: **(1)** decrease in richness and diversity (**Fig. S7A**); **(2)** decrease in the amount of OTUs (**Fig. S7B**); and **(3)** increase in dominance (α diversity) (**Fig. S7C**). In addition, the change in pH resulted in a population turnover (β diversity) (Permanova test, *p* < 0.05, **Table S7**). The effect of temperature, however, was much less pronounced (**Fig. S8A-F**), but was significant at a pH of 6.5 in regards to population turnover (Permanova test, *p* < 0.05, **Table S8**), and most pronounced between samples that were collected at 44°C and 50°C at a pH of 6.5 (Permanova test, *p* < 0.05, **Table S9**), while it was not significant at the pH of 5.0 (**Table S8**).

With the environmental conditions of pH and temperature, we are able to steer the microbiome to a dominant *Lactobacillus* spp. to result in a maximum D-lactate yield (at a pH of 5.0 and a temperature of 50°C), a maximum galactose consumption rate (at a pH of 5.0 and a temperature of 44°C), and a minimum side-product yield (at a pH of 5.0 and a temperature of 44°C). The temperature did affect the kinetics by utilizing the optimum growth conditions for *Lactobacillus* spp. because we observed the highest volumetric D-lactate production rates at a temperature of 44°C. At a pH of 5.0, the operator would need to choose between the maximum volumetric D-lactate production rate and volumetric galactose consumption rate with a minimum of side products at 44°C, or the maximum D-lactate yield and exclusivity at 50°C. Thus, if the most critical performance parameter is D-lactate purity, we found that the reactor microbiome should be operated at a pH of 5.0 and a temperature of 47-50°C to produce D-lactate exclusively.

## 4. Conclusion

- A lactose-rich waste stream from the dairy industry, such as acid whey, was exclusively converted to D-lactate in a continuously operated bioreactor with a reactor microbiome. The process was started *via* autoinoculation. Thus, an external inoculum was avoided.
- D-lactate production required a mildly acidic pH of 5.0 and temperatures ≥ 44°C. High D-lactate yields and volumetric production rates correlated with dominant *Lactobacillus* spp. (> 95% of relative abundance).
- A switch to exclusive L-lactate production was not achieved. A shift from D-lactate towards DL-lactate production required a near-neutral pH of 6.5. L-lactate production correlated with a high relative abundance of *Streptococcus* spp. and *Enterococcus* spp.
- The pH was a powerful process parameter to steer the lactate-isomer yields by affecting microbiome dynamics, while temperature adjustments improved bioprocess kinetics.

## Data availability

Data will be made available on request.

## Acknowledgments

This work was supported by the BMBF VIP + Program (Award # 1336186), and through the Alexander von Humboldt Foundation in the framework of the Alexander von Humboldt Professorship, which was awarded to LTA. The authors wish to thank Heike Budde for performing MiSeq Illumina sequencing (Max Planck Institute for Biology, Tübingen – Germany). In addition, the workers at Friesland-Campina are thanked for providing acid whey.

## Supplementary materials

Supplementary material associated with this article can be found, in the online version, at

