## Supporting Materials for "Exclusive D-lactate-isomer production during a reactor-microbiome conversion of lactose-rich waste by controlling pH and temperature"

### S 1 Material and methods

#### S 1.1 Composition of the acid-whey batch and the acid whey in the feed tank

**Table S1.** Composition of the acid-whey batch upon arrival. Reported values are mean values, and the errors represent the standard deviation (SD) between measurements (n = 3 for all values, except for the COD and TOC measurements with n = 2).

| Collection date | 21/09/2021 |
|---|---|
| Collection site | FrieslandCampina (Cologne, Germany) |
| pH | 4.29 ± 0.01 |
| TS | 56.6 ± 1.57 |
| VS | 49.1 ± 1.55 |
| TSS | 0.07 ± 0.00 |
| VSS | 0.07 ± 0.00 |
| TCOD (g COD $L^{-1}$) | 66.0 ± 1.50 |
| SCOD (g COD $L^{-1}$) | 65.3 ± 1.35 |
| TOC (g C $L^{-1}$) | 23.7 ± 0.13 |
| DOC (g C $L^{-1}$) | 23.4 ± 0.04 |
| SCOD-to-DOC ratio | 2.80 |
| Lactose (mmol $L^{-1}$) | 109 ± 0.85 |
| Galactose (mmol $L^{-1}$) | 35.2 ± 0.35 |
| Lactate (mmol $L^{-1}$) | 86.6 ± 0.80 |
| Calculated SCOD* (g COD $L^{-1}$) | 56.8 ± 0.47 |
| Calculated SCOD*-to-SCOD ratio | 0.87 |
| Calculated DOC** (g C $L^{-1}$) | 21.3 ± 0.18 |
| Calculated DOC**-to-DOC ratio | 0.91 |
| Calculated SCOD*-to-DOC** ratio | 2.66 |

* COD conversions were: 0.384, 0.192, and 0.096 g COD $mmol^{-1}$ for lactose, galactose, and lactate, respectively.
** DOC conversions were 0.144, 0.072, and 0.036 g C $mmol^{-1}$ for lactose, galactose, and lactate, respectively. The total calculated SCOD and DOC include only lactose, galactose, and lactic acid, which are the predominant sources of SCOD and DOC in acid whey.

**Table S2.** Compound concentration in the acid-whey substrate that was measured during P-I, P-II, and P-III. Reported values are mean values, and the errors represent the standard deviation between measurements (n = 28).

| Substrate | Concentration [mmol $L^{-1}$] |
|---|---|
| **Lactose** | 112 ± 23.1 |
| **Galactose** | 31.9 ± 7.25 |
| **Glucose** | 9.14 ± 1.80 |
| **Lactate*** | 83.1 ± 16.8 |
| **D-Lactate**** | 1.75 ± 1.79 |
| **L-Lactate**** | 88.4 ± 17.7 |
| **Ethanol** | 7.66 ± 7.34 |

* numbers based on HPLC measurement (see section 2.4 in the main manuscript)
** numbers based on D-/L-lactate (rapid) assay kit (see section 2.4 in the main manuscript)

#### S 1.2 Bioreactor system

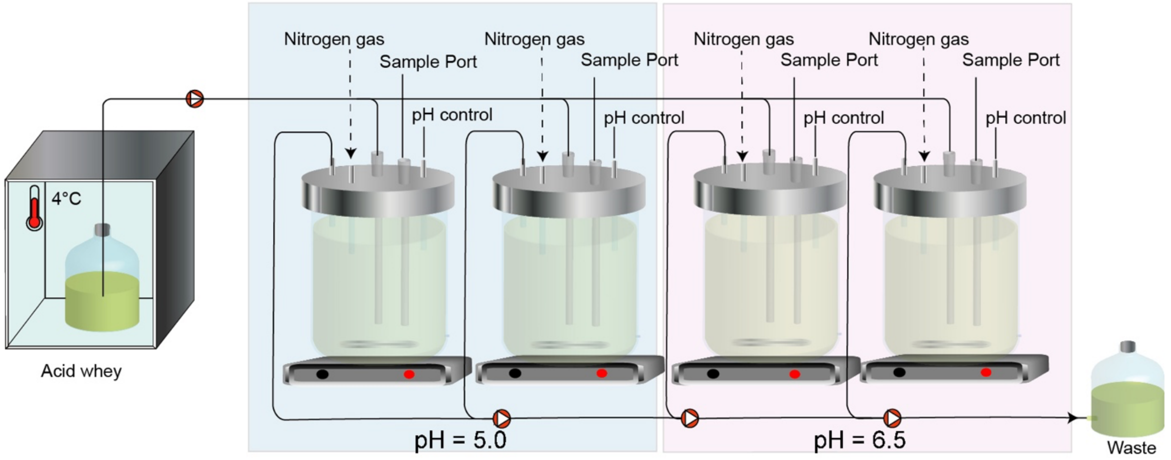

**Figure S1.** Flow scheme of the bioreactor system.

**Table S3.** Working volumes and pH level of reactor 1-4 that were used in the experimental setup.

| | Reactor 1 | Reactor 2 | Reactor 3 | Reactor 4 |
|---|---|---|---|---|
| **pH [-]** | 5.0 | | 6.5 | |
| **Working volume [mL]** | 250 mL | 265 mL | 246 mL | 266 mL |

#### S 1.3 Microbial community analysis

**Table S4.** Primer sequences used to amplify the V4 region of the 16S rRNA gene.

| Forward overhang | Forward primer linker | 515F forward primer |
|---|---|---|
| TCGTCGGCAGCGTCAGATGTGTATAAGAGACAG | GT | GTGYCAGCMGCCGCGGTAA |
| **Reverse overhang** | **Reverse primer linker** | **806R reverse primer** |
| GTCTCGTGGGCTCGGAGATGTGTATAAGAGACAG | CC | GGACTACNVGGGTWTCTAAT |

#### S 1.4 Microscopy

Samples for phase-contrast microscopy were taken during P-II on Day 40 and during P-III on Day 55 to assess differences in the microbial composition of the reactor duplicates based on cell morphology. We centrifuged 1 mL of the liquid sample at 16,740 × g for 6 min at room temperature. The resulting supernatant was discarded, and the pelleted biomass was resuspended in 500 µL of 0.01 x PBS. 2 µL of the cell suspension was transferred onto a microscopy slide. Microscopy was performed with an Olympus BX41 light microscope (Olympus, Tokyo, Japan). Photographs were taken with a 100-fold magnification objective using a camera (Blackfly S USB 3, FLIR Systems, Wilsonville, OR) and the Micro-Manager software (version: 2.0.0-beta3). Image editing and analysis were performed with ImageJ v 1.53f51 (Rasband, W.S., ImageJ, U. S. National Institutes of Health, Bethesda, Maryland, USA).

#### S 1.5 Data processing and calculations

**Volumetric LGGlu loading rate** $[mmol\ C\ L^{-1} d^{-1}]$

$$\frac{c_{L,feed} \times 12 + c_{G,feed} \times 6 + c_{Glu,feed} \times 6}{HRT} \quad \textbf{Eq. S1}$$

Where:

$c_{x,feed}$ concentration of compound $x$ in the acid-whey feed in [mmol L$^{-1}$]
L: lactose; G: Galactose, Glu: Glucose

$HRT$ hydraulic retention time in [d]

**Volumetric LGGlu consumption rate** $[mmol\ C\ L^{-1} d^{-1}]$

$$\frac{(c_{L,feed} - c_L) \times 12 + (c_{G,feed} - c_G) \times 6 + (c_{Glu,feed} - c_{Glu}) \times 6}{HRT} \quad \textbf{Eq. S2}$$

Where:

$c_{x,feed}$ concentration of compound $x$ in the acid-whey feed in [mmol L$^{-1}$]

$c_x$ concentration of compound $x$ in the fermentation broth in [mmol L$^{-1}$]
L: lactose; G: Galactose, Glu: Glucose

$HRT$ hydraulic retention time in [d]

**Volumetric net production rate $r_x$ of specific product $x$** $[mmolC\ L^{-1} d^{-1}]$

$$\frac{(c_x - c_{x,feed}) \times N_{C,x}}{HRT} \quad \textbf{Eq. S3}$$

Where:

$c_x$ concentration of compound $\boldsymbol{x}$ in the fermentation broth [mmol L$^{-1}$]

$c_{x,feed}$ concentration of compound $\boldsymbol{x}$ in the acid-whey feed in [mmol L$^{-1}$]

$N_{C,x}$ number of carbon atoms per molecule of compound $x$

Please note that rates for lactate (isomer) production are reported as net lactate (isomer) production rate within the bioreactor system and have been corrected for the influent lactate (isomers) feeding rate, which is inherent to feeding acid whey. Negative volumetric net production rates are considered as volumetric consumption rates.

**Product yield of specific product $x$** $[mmol\ C\ mmol\ C^{-1}]$

$$\frac{r_x}{\text{Volumetric LGGlu consumption rate}} \quad \textbf{Eq. S4}$$

### S 2 Results

#### S 2.1 16S rRNA gene sequencing quality using *Illumina* MiSeq

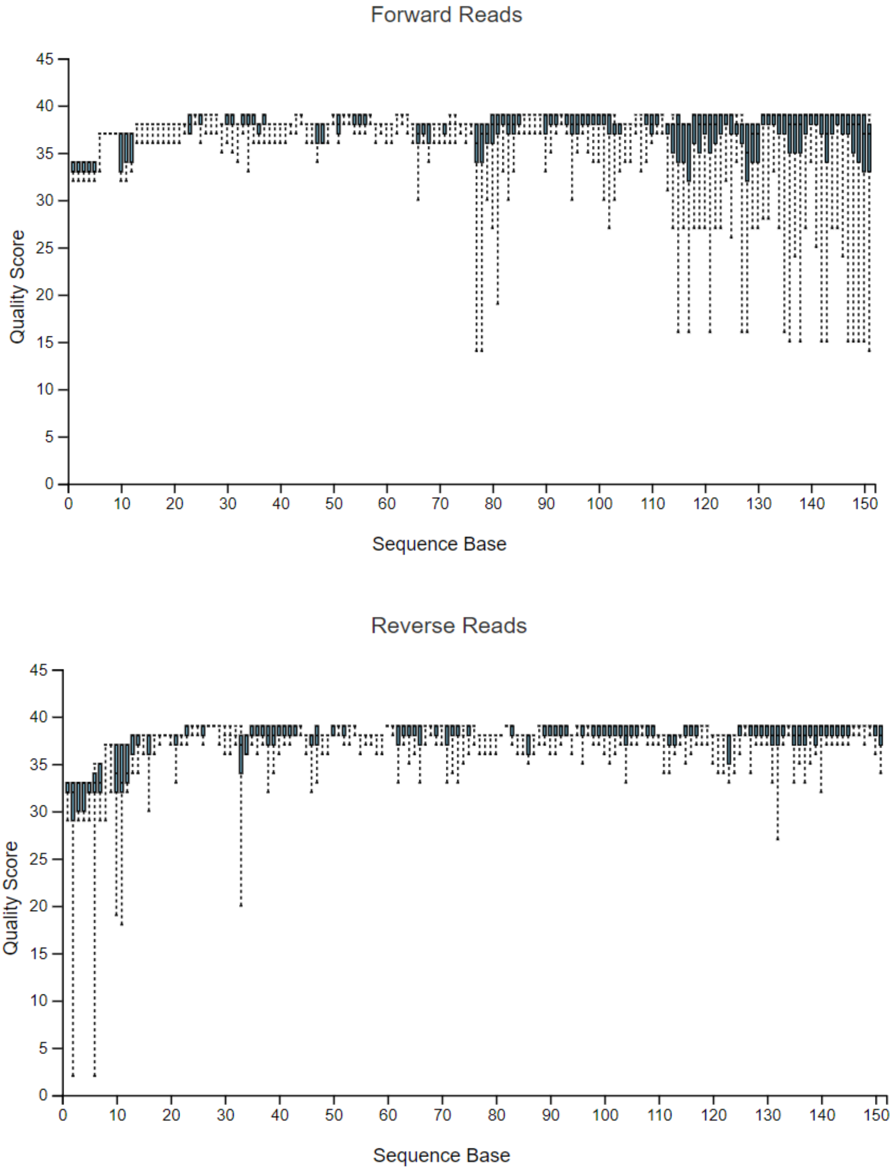

**Figure S2.** Quality scores of Illumina 16S rRNA gene sequence reads: (top) forward sequencing reads; and (bottom) reverse sequencing reads.

According to *Illumina*® protocol, all sequence reads were high quality, and sequencing revealed 415 operational taxonomic units (OTUs) among all collected samples. The median sequence count was 386,707 for forward and reverse reads. The number of sequences per sample varied from a minimum of 312,877 to a maximum of 479,725. We identified 13 OTUs that comprised at least 1% of the relative abundance at a genus-level annotation. It is important to note that the freshly thawed acid-whey samples (see section 2.1 in the main manuscript) have been sequenced on a different *Illumina* sequencing run (sequence counts of 64,416 and 78,106 for both samples, respectively).

#### S 2.2 Compound concentration *vs.* time – complete time-course data

##### S 2.2.1 Mildly-acidic pH of 5.0

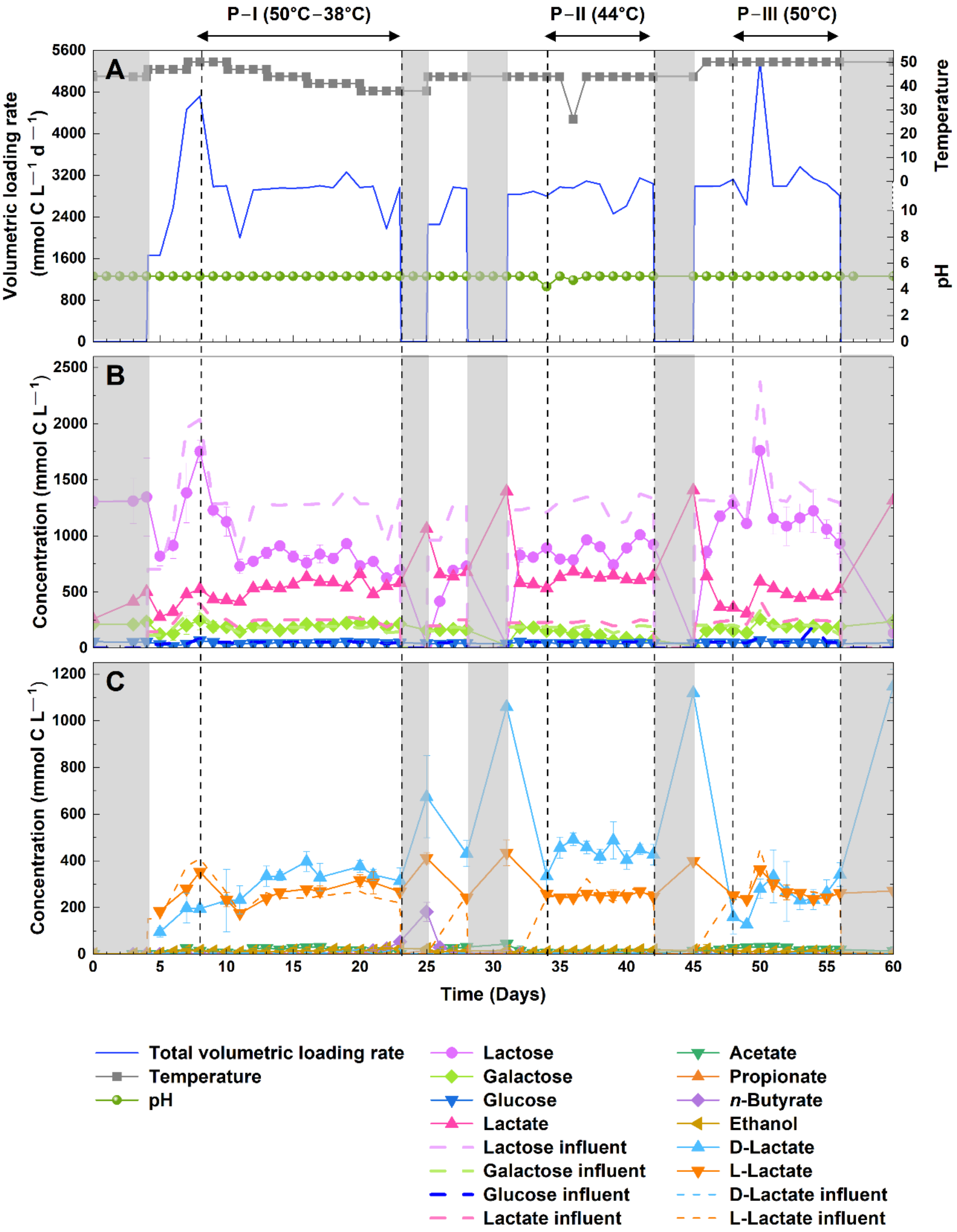

**Figure S3**. Performance of the reactor duplicates operated at pH 5 throughout the operating period: **A)** line plots of operating parameters: total volumetric loading rate (*i.e.*, lactose, galactose, glucose, and lactate), temperature, and pH**; B)** concentrations of lactose, galactose, glucose, and lactate in the influent and fermentation broth; and **C)** concentrations of D-lactate, L-lactate (influent and fermentation broth), and side products (*i.e.*, acetate, propionate, *n*-butyrate, and ethanol). Solid lines represent compound concentration in the fermentation broth while dashed lines represent compound concentration in the acid-whey feed. Error bars represent the standard deviations among duplicate reactors.

##### S 2.2.2 Near-neutral pH of 6.5

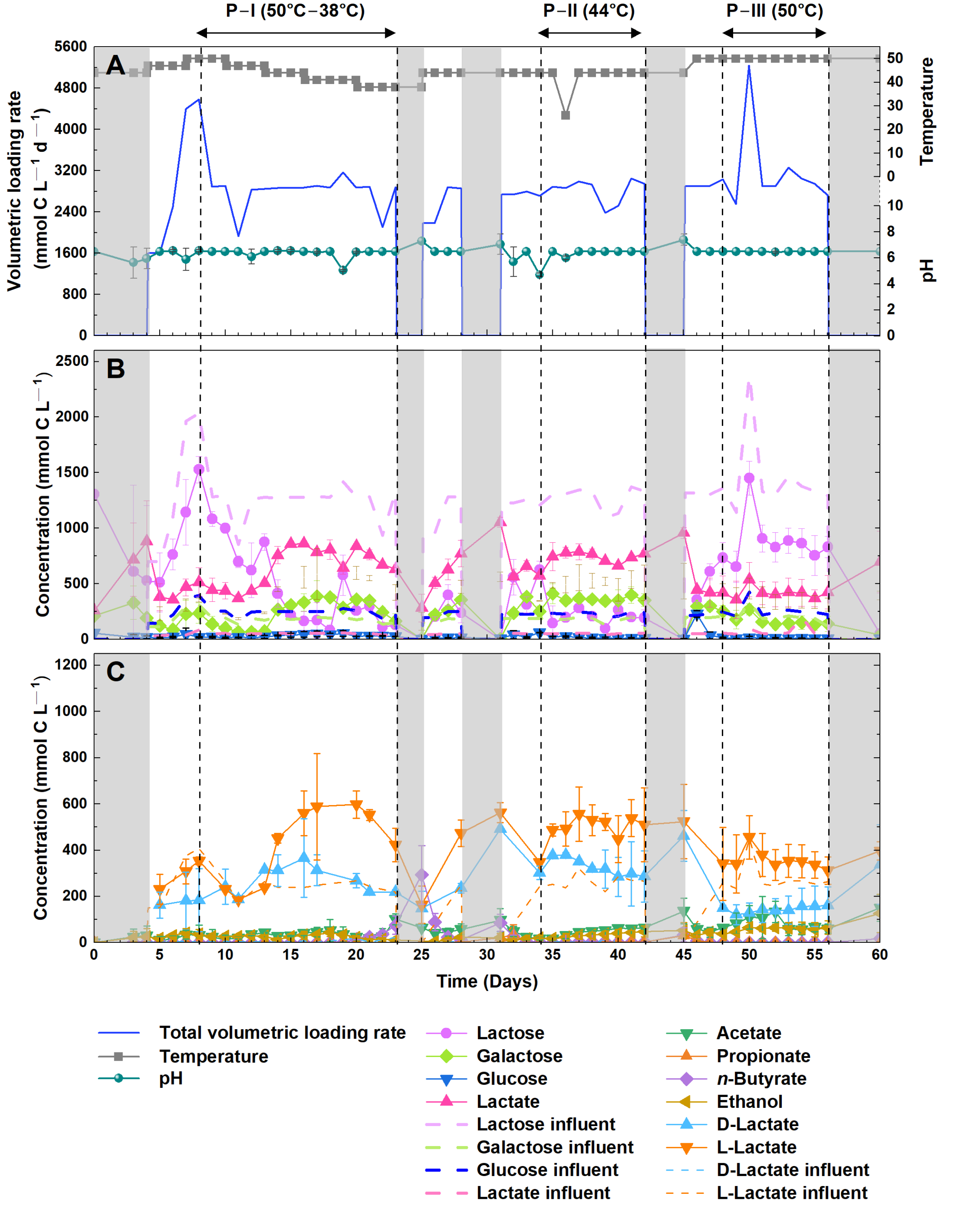

**Figure S4.** Performance of the reactor duplicates operated at pH 6.5 throughout the operating period: **A)** line plots of operating parameters: total volumetric loading rate (*i.e.*, lactose, galactose, glucose, and lactate), temperature, and pH**; B)** concentrations of lactose, galactose, glucose, and lactate in the influent and fermentation broth; and **C)** concentrations of D-lactate, L-lactate (influent and fermentation broth), and side products (*i.e.*, acetate, propionate, *n*-butyrate, and ethanol). Solid lines represent compound concentration in the fermentation broth while dashed lines represent compound concentration in the acid-whey feed. Error bars represent the standard deviations among duplicate reactors.

**Table S5.** Overview of kinetic bioprocess performance parameters at steady state during P-I.

| Parameter | | pH 5.0 | | | | | pH 6.5 | | | | |
|---|---|---|---|---|---|---|---|---|---|---|---|
| Temperature[°C] | | 50 | 47 | 44 | 41 | 38 | 50 | 47 | 44 | 41 | 38 |
| **Lactose-consumption rate [mmol C L$^{-1}$ d$^{-1}$]** | Mean ± SD | 334 ± 193 (n = 5) | 579 ± 297 (n = 6) | 752 ± 139 (n = 6) | 822 ± 88.7 (n = 8) | 809 ± 246 (n = 6) | 557 ± 260 (n = 6) | 664 ± 409 (n = 6) | 169x10$^{1}$ ± 229 (n = 6) | 174x10$^{1}$ ± 283 (n = 8) | 167x10$^{1}$ ± 291 (n = 6) |
| | Median | 362 | 721 | 758 | 838 | 853 | 486 | 673 | 176x10$^{1}$ | 171x10$^{1}$ | 164x10$^{1}$ |
| | Skewness | 0.23 | -0.94 | -0.1 | -0.81 | -0.39 | 0.92 | 0.9 | -1.08 | -0.91 | -0.18 |
| **Galactose-consumption rate [mmol C L$^{-1}$ d$^{-1}$]** | Mean ± SD | 62.4 ± 84.4 (n = 5) | 0.22 ± 0.55 (n = 6) | 22.0 ± 27.4 (n = 6) | 3.01 ± 8.53 (n = 8) | 0.00 ± 0.00 (n = 6) | 123 ± 92.2 (n = 6) | 163 ± 60.4 (n = 6) | 0.00 ± 0.00 (n = 6) | 0.00 ± 0.00 (n = 8) | 7.23 ± 17.7 (n = 6) |
| | Median | 45.6 | 0.00 | 11.6 | 0.00 | 0.00 | 150 | 168 | 0.00 | 0.00 | 0.00 |
| | Skewness | 1.68 | 2.45 | 0.8 | 2.83 | -- | -0.41 | -0.8 | -- | -- | 2.45 |
| **Glucose-consumption rate [mmol C L$^{-1}$ d$^{-1}$]** | Mean ± SD | 33.5 ± 20.8 (n = 5) | 12.1 ± 7.66 (n = 6) | 12.3 ± 2.99 (n = 15) | 8.74 ± 6.40 (n = 8) | 6.23 ± 5.47 (n = 6) | 75.3 ± 27.8 (n = 6) | 47.2 ± 21.4 (n = 6) | 24.4 ± 19.5 (n = 6) | 24.5 ± 19.1 (n = 8) | 31.7 ± 14.3 (n = 6) |
| | Median | 29.8 | 13.4 | 13.1 | 8.92 | 6.87 | 66.4 | 36.1 | 30.8 | 37.6 | 38.0 |
| | Skewness | -0.65 | -0.37 | -0.67 | 0.11 | -0.2 | 0.48 | 1.12 | -0.63 | -0.66 | -1.02 |
| **Lactate production rate [mmol C L$^{-1}$ d$^{-1}$]** | Mean ± SD | 292 ± 147 (n = 5) | 461 ± 73.9 (n = 6) | 543 ± 79.0 (n = 6) | 557 ± 95.2 (n = 8) | 513 ± 171 (n = 6) | 277 ± 149 (n = 6) | 358 ± 59.9 (n = 6) | 960 ± 106 (n = 6) | 847 ± 164 (n = 8) | 765 ± 108 (n = 6) |
| | Median | 236 | 477 | 524 | 562 | 586 | 288 | 347 | 996 | 894 | 797 |
| | Skewness | 0.28 | -1.87 | 0.63 | -0.23 | -1.86 | -0.51 | 0.51 | -1.83 | -1.02 | -0.42 |
| **D-lactate production rate [mmol C L$^{-1}$ d$^{-1}$]** | Mean ± SD | 347 ± 134 (n = 4) | 467 ± 120 (n = 4) | 603 ± 70.4 (n = 4) | 571 ± 68.2 (n = 4) | 525 ± 74.9 (n = 4) | 348 ± 160 (n = 4) | 413 ± 122 (n = 4) | 559 ± 182 (n = 4) | 467 ± 93.1 (n = 4) | 346 ± 23.0 (n = 4) |
| | Median | 320 | 477 | 589 | 590 | 544 | 383 | 419 | 519 | 446 | 352 |
| | Skewness | 1.11 | -0.47 | 1.03 | -1.19 | -1.34 | -0.8 | -0.07 | 0.88 | 1.27 | -0.94 |
| **L-lactate production rate [mmol C L$^{-1}$ d$^{-1}$]** | Mean ± SD | 0.00 ± 0.00 (n = 4) | 2.34 ± 4.67 (n = 4) | 53.8 ± 23.8 (n = 4) | 57.3 ± 42.8 (n = 4) | 92.8 ± 49.0 (n = 4) | 0.00 ± 0.00 (n = 4) | 8.27 ± 13.0 (n = 4) | 444 ± 141 (n = 4) | 559 ± 229 (n = 4) | 428 ± 125 (n = 4) |
| | Median | 0.00 | 0.00 | 53.9 | 56.1 | 79.8 | 0.00 | 2.88 | 399 | 551 | 457 |
| | Skewness | -- | 2 | -0.04 | 0.17 | 1.4 | -- | 1.76 | 1.61 | 0.19 | -1.16 |
| **L-lactate consumption rate [mmol C L$^{-1}$ d$^{-1}$]** | Mean ± SD | 72.0 ± 39.0 (n = 4) | 23.55 ± 23.1 (n = 4) | 0.00 ± 0.00 (n = 4) | 0.00 ± 0.00 (n = 4) | 0.00 ± 0.00 (n = 4) | 71.7 ± 28.0 (n = 4) | 21.0 ± 25.6 (n = 4) | 0.00 ± 0.00 (n = 4) | 0.00 ± 0.00 (n = 4) | 0.00 ± 0.00 (n = 4) |
| | Median | 87.0 | 20.1 | 0.00 | 0.00 | 0.00 | 64.0 | 16.1 | 0.00 | 0.00 | 0.00 |
| | Skewness | 1.85 | -0.75 | -- | -- | -- | -1.39 | -0.48 | -- | -- | -- |
| **Lactate yield [mmol C mmol C$^{-1}$]** | Mean ± SD | 0.84 ± 0.64 (n = 5) | 1.10 ± 0.76 (n = 6) | 0.69 ± 0.04 (n = 6) | 0.67 ± 0.09 (n = 8) | 0.71 ± 0.40 (n = 6) | 0.43 ± 0.29 (n = 6) | 0.52 ± 0.27 (n = 6) | 0.56 ± 0.02 (n = 6) | 0.48 ± 0.06 (n = 8) | 0.47 ± 0.13 (n = 6) |
| | Median | 0.71 | 0.68 | 0.70 | 0.70 | 0.64 | 0.38 | 0.48 | 0.56 | 0.47 | 0.50 |
| | Skewness | 1.61 | 1.26 | -1.33 | -1.15 | 0.26 | 0.63 | 0.53 | 0.1 | 0.23 | -0.4 |
| **D-lactate yield [mmol C mmol C$^{-1}$]** | Mean ± SD | 0.82 ± 0.35 (n = 5) | 1.35 ± 0.76 (n = 4) | 0.78 ± 0.05 (n = 4) | 0.67 ± 0.05 (n = 4) | 0.55 ± 0.12 (n = 4) | 0.42 ± 0.24 (n = 4) | 0.67 ± 0.11 (n = 4) | 0.33 ± 0.08 (n = 4) | 0.26 ± 0.03 (n = 4) | 0.19 ± 0.03 (n = 4) |
| | Median | 0.78 | 1.17 | 0.78 | 0.69 | 0.58 | 0.43 | 0.66 | 0.35 | 0.26 | 0.19 |
| | Skewness | 0.26 | 0.77 | 0.29 | -1.75 | -0.52 | -0.17 | 0.38 | -1.16 | -0.28 | -0.03 |
| **L-lactate yield [mmol C mmol C$^{-1}$]** | Mean ± SD | 0.00 ± 0.00 (n = 4) | 0.01 ± 0.02 (n = 4) | 0.07 ± 0.03 (n = 4) | 0.07 ± 0.05 (n = 4) | 0.10 ± 0.07 (n = 4) | 0.00 ± 0.00 (n = 4) | 0.02 ± 0.03 (n = 4) | 0.26 ± 0.06 (n = 4) | 0.32 ± 0.15 (n = 4) | 0.24 ± 0.09 (n = 4) |
| | Median | 0.00 | 0.00 | 0.07 | 0.07 | 0.08 | 0.00 | 0.01 | 0.24 | 0.31 | 0.25 |
| | Skewness | -- | 2 | -0.13 | -0.51 | 1.85 | -- | 1.76 | 1.71 | 0.06 | -0.73 |
| **Side-product yield [mmol C mmol C$^{-1}$]** | Mean ± SD | 0.06 ± 0.03 (n = 5) | 0.07 ± 0.02 (n = 6) | 0.06 ± 0.01 (n = 6) | 0.05 ± 0.03 (n = 8) | 0.10 ± 0.03 (n = 6) | 0.09 ± 0.08 (n = 6) | 0.10 ± 0.05 (n = 6) | 0.04 ± 0.01 (n = 6) | 0.05 ± 0.02 (n = 8) | 0.09 ± 0.05 (n = 6) |
| | Median | 0.06 | 0.06 | 0.06 | 0.04 | 0.09 | 0.06 | 0.09 | 0.04 | 0.05 | 0.08 |
| | Skewness | -0.63 | 1.99 | -0.52 | 0.67 | 7 | 1.29 | 1.33 | 0.66 | 0.58 | 0.98 |

| Parameter | | [illegible] | | [illegible] | |
|---|---|---|---|---|---|
| **pH** | | 5.0 | 6.5 | 5.0 | 6.5 |
| **Lactose-consumption rate [mmol C L$^{-1}$ d$^{-1}$]** | Mean ± SD | 657 ± 149<br>(n = 18) | 171x10$^{1}$ ± 322<br>(n = 18) | 434 ± 311<br>(n = 17) | 943 ± 294<br>(n = 18) |
| | Median | 651 | 181x10$^{1}$ | 456 | 887 |
| | Skewness | 0.02 | -1.6 | 0.52 | 1.12 |
| **Galactose-consumption rate [mmol C L$^{-1}$ d$^{-1}$]** | Mean ± SD | 125 ± 76.0<br>(n = 18) | 0.00 ± 0.00<br>(n = 18) | 36.9 ± 48.9<br>(n = 17) | 66.0 ± 59.4<br>(n = 18) |
| | Median | 114 | 0.00 | 12.4 | 57.4 |
| | Skewness | 0.55 | -- | 1.41 | 0.46 |
| **Glucose-consumption rate [mmol C L$^{-1}$ d$^{-1}$]** | Mean ± SD | 15.0 ± 3.71<br>(n = 18) | 54.4 ± 23.7<br>(n = 18) | 13.8 ± 11.7<br>(n = 17) | 76.6 ± 21.0<br>(n = 18) |
| | Median | 15.0 | 61.4 | 9.9 | 70.3 |
| | Skewness | -0.09 | -1.36 | 1.84 | 2.47 |
| **Lactate production rate [mmol C L$^{-1}$ d$^{-1}$]** | Mean ± SD | 664 ± 88.5<br>(n = 18) | 828 ± 123<br>(n = 18) | 356 ± 138<br>(n = 17) | 270 ± 167<br>(n = 18) |
| | Median | 659 | 839 | 356 | 252 |
| | Skewness | 0.05 | -0.71 | 0.39 | -0.03 |
| **D-lactate production rate [mmol C L$^{-1}$ d$^{-1}$]** | Mean ± SD | 722 ± 94.6<br>(n = 18) | 535 ± 106<br>(n = 18) | 420 ± 136<br>(n = 17) | 235 ± 76.8<br>(n = 18) |
| | Median | 723 | 570 | 419 | 242 |
| | Skewness | -0.62 | -0.9 | 0.13 | 0.3 |
| **L-lactate production rate [mmol C L$^{-1}$ d$^{-1}$]** | Mean ± SD | 10.5 ± 21.2<br>(n = 18) | 385 ± 144<br>(n = 18) | 5.42 ± 18.5<br>(n = 13) | 140 ± 114<br>(n = 18) |
| | Median | 0.00 | 382 | 0.00 | 110 |
| | Skewness | 2.67 | -0.26 | 3.59 | 0.43 |
| **L-lactate consumption rate [mmol C L$^{-1}$ d$^{-1}$]** | Mean ± SD | 28.1 ± 36.3<br>(n = 18) | 0.00 ± 0.00<br>(n = 18) | 16.4 ± 19.4<br>(n = 13) | 8.71 ± 26.7<br>(n = 18) |
| | Median | 16.5 | 0.00 | 7.91 | 0.00 |
| | Skewness | -1.5 | -- | -1.01 | -3.18 |
| **Lactate yield [mmol C mmol C$^{-1}$]** | Mean ± SD | 0.86 ± 0.17<br>(n = 18) | 0.48 ± 0.07<br>(n = 18) | 0.81 ± 0.36<br>(n = 15) | 0.25 ± 0.15<br>(n = 18) |
| | Median | 0.78 | 0.48 | 0.80 | 0.22 |
| | Skewness | 1.28 | -0.07 | 0.13 | 0.23 |
| **D-lactate yield [mmol C mmol C$^{-1}$]** | Mean ± SD | 0.93 ± 0.15<br>(n = 18) | 0.32 ± 0.10<br>(n = 18) | 0.97 ± 0.38<br>(n = 15) | 0.22 ± 0.07<br>(n = 18) |
| | Median | 0.85 | 0.32 | 0.93 | 0.22 |
| | Skewness | 0.88 | 0.65 | 0.19 | 0.18 |
| **L-lactate yield [mmol C mmol C$^{-1}$]** | Mean ± SD | 0.01 ± 0.03<br>(n = 18) | 0.22 ± 0.06<br>(n = 18) | 0.01 ± 0.03<br>(n = 13) | 0.13 ± 0.10<br>(n = 18) |
| | Median | 0.00 | 0.20 | 0.00 | 0.11 |
| | Skewness | 2.42 | -0.19 | 3.53 | 0.59 |
| **Side-product yield [mmol C mmol C$^{-1}$]** | Mean ± SD | 0.02 ± 0.01<br>(n = 18) | 0.06 ± 0.02<br>(n = 18) | 0.11 ± 0.09<br>(n = 15) | 0.15 ± 0.05<br>(n = 16) |
| | Median | 0.02 | 0.06 | 0.07 | 0.14 |
| | Skewness | 0.9 | 0.01 | 1.27 | 1.11 |

#### S 2.4 Volatile suspended solids during P-II and P-III

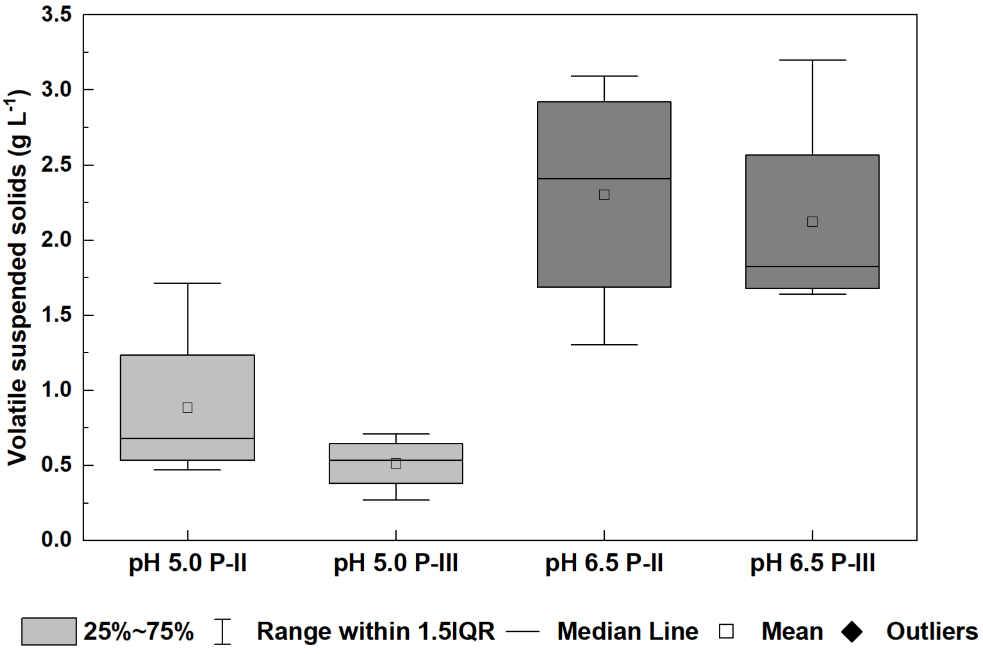

**Figure S5.** Volatile suspended solids measurements of the fermentation broth during P-II and P-III at a mildly-acid pH of 5.0 and near-neutral pH of 6.5. Boxplot: box: $Q_{0.25}$–$Q_{0.75}$; band inside the box: $Q_{0.5}$ (median); end of the whiskers:1.5 × interquartile range; square: mean; (n = 4 ).

#### S 2.5 Phase-contrast microscopy

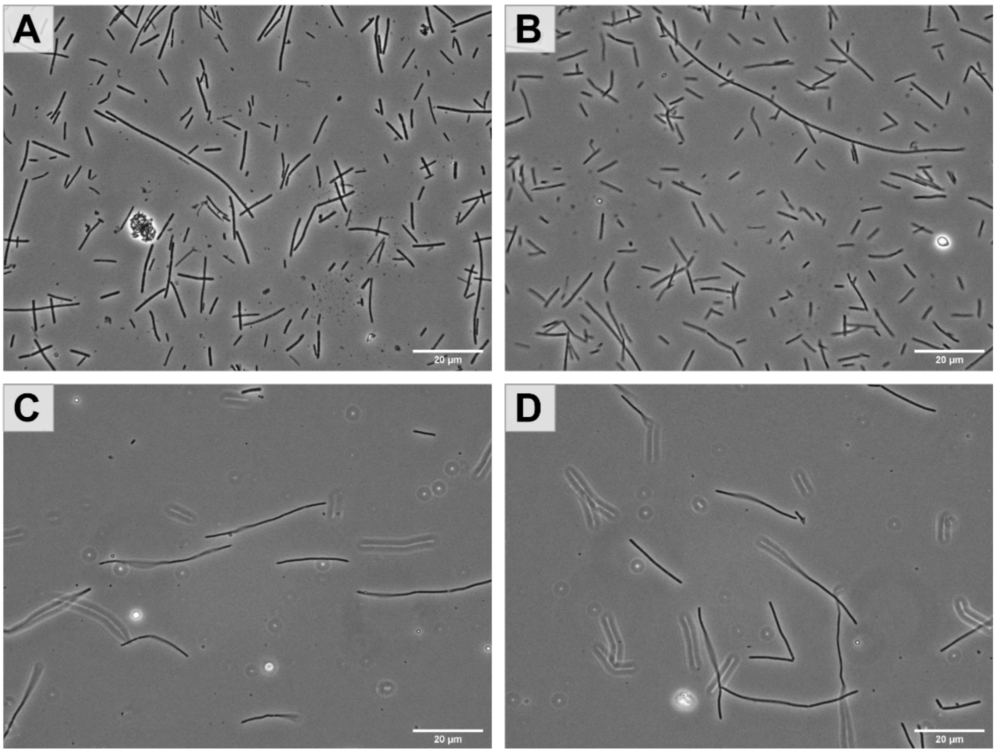

**Figure S6**. Phase-contrast micrographs of bioreactor samples: **A)** pH 5.0 and 44°C for reactor 1 (P-II); **B)** pH 5.0 and 44°C for reactor 2 (P-II); **C)** pH 5.0 and 50°C for reactor 1 (P-III); and **D)** pH 5.0 and 50°C for reactor 2 (P-III). The scale bar represents 20 µm.

#### S 2.6 α diversity and β diversity metrics

##### S 2.6.1 α diversity for pH 5.0 *vs.* pH 6.5

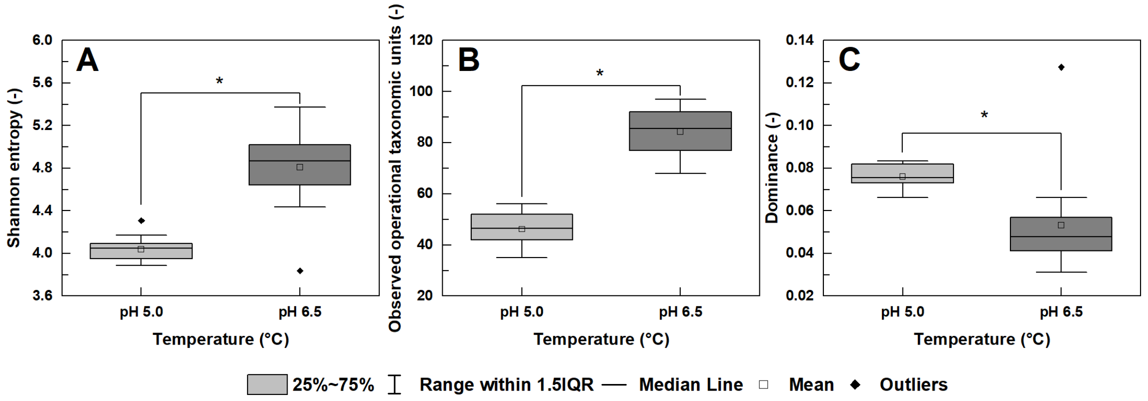

**Figure S7.** α-Diversity metrics for pH 5.0 *vs.* pH 6.5 using pairwise Kruskal-Wallis test that was calculated by QIIME2: **A)** Shannon entropy; **B)** OTUs; and **C)** dominance; n = 14 per group; * = $p< 0.01$.

##### S 2.6.2 α diversity between temperature groups

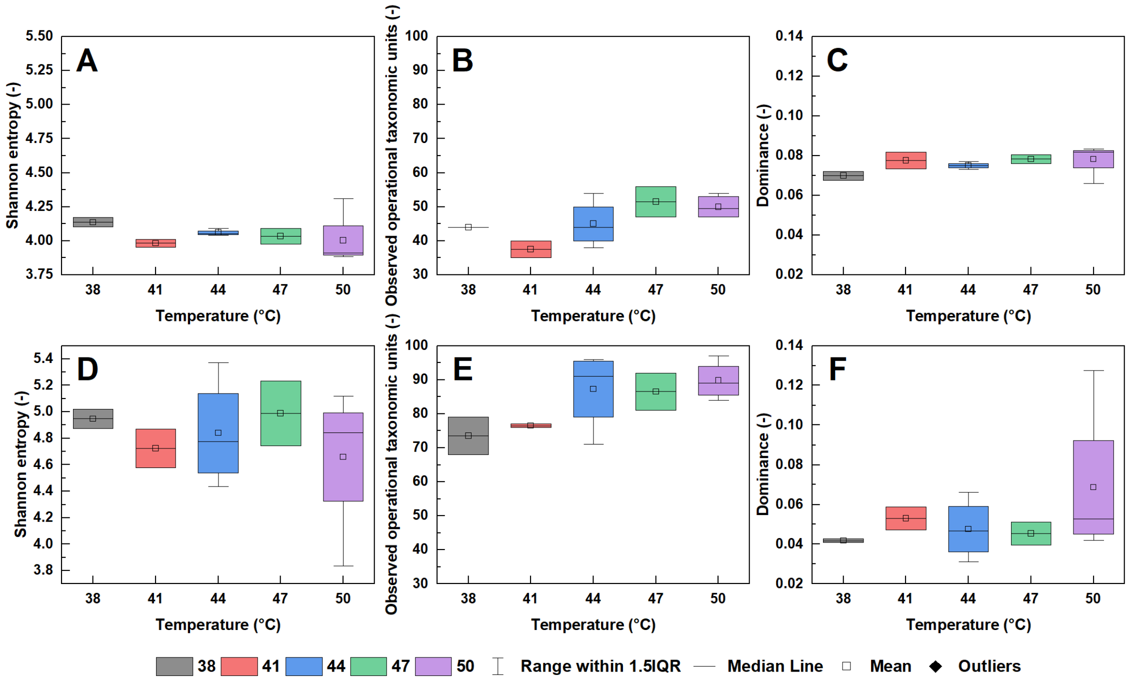

**Figure S8.** α-Diversity metrics between temperature groups on pairwise Kruskal-Wallis that was calculated by QIIME2: **A)** Shannon entropy at a pH of 5.0; **B)** OTUs at a pH of 5.0; **C)** Dominance at a pH of 5.0; **D)** Shannon entropy at a pH of 6.5; **E)** OTUs at a pH of6.5; and **F)** Dominance at a pH of 6.5. n = 2 for 38°C (grey), 41°C (red), and 47°C (green), while n = 4 for 44°C (blue) and 50°C (purple).

#### S 2.6.3 β diversity for pH 5.0 *vs.* pH 6.5

**Table S7.** Statistical differences in β diversity pH 5.0 *vs.* pH 6.5 based on the Permanova test calculated by QIIME2.

| Distance Matrix | Sample size | Pseudo-F value | *p*-value |
|---|---|---|---|
| **Bray-Curtis** | 28 | 19.06 | **0.001** |
| **Weighted UniFrac** | 28 | 20.5 | **0.001** |

#### S 2.6.4 β diversity between temperatures at pH 5.0 and pH 6.5

**Table S8.** Statistical differences in β diversity between temperature conditions at pH 5.0 and pH 6.5 based on the Permanova test calculated by QIIME2.

| pH | Distance matrix | Sample size | Pseudo-F value | *p*-value |
|---|---|---|---|---|
| **5** | Bray-Curtis | 14 | 2.76 | **0.046** |
| | Weighted UniFrac | 14 | 1.31 | **0.204** |
| **6.5** | Bray-Curtis | 14 | 5.2 | **0.001** |
| | Weighted UniFrac | 14 | 7.23 | **0.001** |

**Table S9.** Statistical differences in β diversity between statistically significant temperature conditions (44°C *vs.* 50°C) at pH 6.5 based on the Permanova test calculated by QIIME2.

| pH | Distance matrix | Group 1 | Group 2 | Sample size | Pseudo-F value | *p*-value |
|---|---|---|---|---|---|---|
| **6.5** | Bray-Curtis | 44 | 50 | 8 | 6.73 | **0.02** |
| | Weighted UniFrac | 44 | 50 | 8 | 10.47 | **0.02** |
